# Hematopoietic Stem Cells Modulate Tumor Immune-Environment to Target Triple-Negative Breast Cancer via Altering Mitochondrial Bioenergetics

**DOI:** 10.1101/2025.05.07.652451

**Authors:** Sumit Mallick, Siddhartha Biswas, Sudheer Shenoy P, Bipasha Bose

## Abstract

Ontogenic development of Hematopoietic stem cells (HSCs) takes place at diverse anatomical niches. Moreover, during embryonic development, HSCs migrate from the aorta-gonad mesonephros (AGM) to the fetal liver and finally to the bone marrow immediately after childbirth. Hence, the primary residence of adult HSCs in the bone marrow continues replenishing the hematopoietic lineage pool. *In vivo*, the HSC niche is significant and might be harnessed in regenerative medicine. HSCs in various niches have exhibited their respective tropism and proliferations based on the growth factors secreted by the niche. Accordingly, in this work, we hypothesized HSCs tropism towards cancer and stem cell niche of triple-negative breast cancer (TNBC) and HER2^+^, having a high relapse rate for possible cell therapy development. Our results indicate that HSC exclusive tropism towards breast cancer stem cells (CSC) and interaction with the cancer milieu lead to HSC differentiation into T-lymphocyte cells (CD4 & CD8). Moreover, the single-cell type proteomics of the migrated HSCs towards TNBC-CSCs and HER2^+^ cells indicated upregulation of IL-7 and Notch protein and several other upregulated proteins primarily involved in T cell activation and migration-related pathways. Likewise, the metabolomics from HSCs-derived conditioned media-treated CSCs, and HER2^+^ cells showed the capability of HSC-CM in arresting the growth and cell cycle of TNBC-CSC via altered mitochondrial bioenergetics. Hence, this study paves the way toward harnessing the potential of both HSCs and HSC-CM for personalized medicine against TNBC CSCs.

**Highlights:**

1. Hematopoietic stem cells exhibited tumor tropism towards triple negative breast cancer stem cells and HER2^+^ cancer cells.
2. Upon migration in the tumor milieu, HSCs differentiated into T cells, targeted CSCs, and modulated the upregulation of interleukin-7 and notch proteins.
3. HSCs-CM inhibited the cell cycle in TNBC-CSCs by downregulating CDK1, disrupting mitochondrial bioenergetics via upregulated Drp1, and inducing CSC apoptosis.
4. Paracrine signaling from HSCs via conditioned media induced metabolic stress and disrupted key metabolic pathways in TNBC-CSCs.

**Summary:** Hematopoietic stem cells (HSCs) are known for their regenerative potential in the blood system. This study reveals the novel therapeutic potential of hematopoietic stem cells (HSCs) in combating triple-negative breast cancer stem cells (TNBC-CSCs). HSCs exhibited tropism towards TNBC-CSCs, themselves differentiated into T cells within the tumor microenvironment, leading to targeted cell killing of TNBC-CSCs. In addition, paracrine signaling from HSC-conditioned media (CM) induced metabolic stress in TNBC-CSCs and disrupted key metabolic pathways. HSC-CM also disrupted mitochondrial dynamics and function, leading to DNA damage and apoptosis. Critically, HSC-CM impaired TNBC-CSC stemness by downregulating stemness genes, leading to the inhibition of 3D spheroid formation. Hence, our study highlights HSCs as a promising therapeutic strategy for targeting TNBC-CSCs via disrupted metabolic homeostasis and self-renewal.

## Introduction

Hematopoietic stem cells (HSCs) are generally considered the backbone of the adult hematopoietic system as they replenish the entire blood system. After the development of the HSCs in the hemogenic endothelium, they are detected in the aorta-gonad-mesonephros (AGM) region. On embryonic day 10.5, HSCs migrate into the blood and maintain the potential for repopulation (1–11). Apart from the remarkable plasticity and regenerative potential within the hematopoietic system, recent advancements have unveiled an intriguing aspect of HSC biology – their inherent tropism towards specific niches within the body (12–16). This tropism behaviour, driven by intricate various cytokines and chemotactic signals and molecular interactions, modulates diverse biological functions, which underscore the versatility of HSCs beyond conventional hematopoiesis (17–20).

In our studies, we hypothesize that HSCs can show tropism towards breast cancer and cancer stem cells, possibly targeting the CSCs and breast cancer as a strategy for targeting early and metastatic breast cancer (21–25). Here, we have used HER2^+^ and triple-negative breast cancer stem cells. Triple-negative breast cancer (TNBC), characterized by its aggressive phenotype and limited treatment options, stands as a prime candidate for exploring the therapeutic potential of HSCs. Central to TNBC’s resilience and therapeutic resistance are cancer stem cells (CSCs), a subpopulation endowed with self-renewal capacity and tumorigenic potential. Against this backdrop, our study delves into the intricate interplay between HSCs and TNBC CSCs, aiming to exploit the tropism properties of HSCs for targeted anti-cancer strategies. Hence, our hypothesis on the tropism behaviour preference of HSCs towards TNBC microenvironments may serve as a foundation for developing innovative therapeutic modalities specifically tailored to eradicate TNBC CSCs. One interesting aspect is that both HSCs and TNBC microenvironments share some common cytokines and chemokines such as TNF-α, CXCR4, CXCL-12, IL-7, SDF-1, and specific matrix metalloproteinases protein (MMP-2, MMP-9). Such cytokines often crosstalk with each other, possibly leading to HSC tropism toward TNBC cells (21–23). Moreover, the tropism characteristic of HSCs will further help to elucidate their use as drug delivery vehicles toward TNBC-CSCs.

Our initial investigations revealed a striking specificity of HSCs towards TNBC CSCs, as evidenced by exclusive tropism behaviour towards cancer cell niches in static co-culture settings. After these findings, we further explored the molecular underpinnings of HSC-TNBC CSCs microenvironment (TME) interactions. Results established that migratory HSCs differentiated into helper and cytotoxic T cells (CD4 & CD8) in the TME. For robust and more detailed analysis, we further performed a multi-omics approach, including migrated single-cell type proteomics, conditioned media metabolomics, and metabolomics of CM-treated TNBC-CSCs and HER2+ cells, which uncovered a cascade of cellular responses within TNBC CSCs, including apoptosis induction, DNA double-stranded breakage (DSB), cell cycle arrest, and disruption of mitochondrial bioenergetics. Hence, these observations highlight the multifaceted nature of HSC-mediated anti-cancer effects, implicating both intrinsic apoptotic pathways and mitochondrial bioenergetics in the demise of TNBC CSCs.

## Material and Methods

A detailed methodology is provided in the supplementary methods file.

### Cell Culture

In this study, cell lines HL-60, MDA-MB-468, MDA-MB-231, MCF-7, and HaCaT were obtained from the National Centre for Cell Sciences in Pune, India. HL-60 Cells were cultured in RPMI media with growth factors (see the Extended file) (Pepro-Tech, ThermoFisher Scientific). HaCaT and breast cancer cells were cultured in DMEM (Gibco Thermo Scientific), which was supplemented with 10% fetal bovine serum (Hi-Media Laboratories, India), 1% Penicillin and Streptomycin, 1% non-essential amino acids (NEAA), and 1% Glutamax (Gibco Thermo Scientific). At 70% confluency, cells were trypsinized, expanded, and cryopreserved.

### Hematopoietic stem cells (HSCs) culture and sorting

HSCs were derived from HL-60 cell lines cultured in RPMI-1640 initially and then switched over to media containing a growth factor cocktail (SCF; 50ng/mL, FLT3; 20ng/mL, IL-6; 50ng/mL, TPO; 10ng/mL) (**Pepro Tech, ThermoFisher**) with the addition of FBS (26). The cells were stained with CD34 and CD45 antibodies (**Extended file 2) at 4°C for 30 minutes**. The cells were then sorted based on CD34 and CD45 expression using a Bio-Rad S3E cell sorter. The sorted cells (CD34^+^ and CD45^+^) were cultured in serum-free media with a growth factor cocktail for further experiments.

### Collection of conditioned media (CM) from sorted HSCs

Sorted 10 × 10^6 HSCs were cultured in RPMI media with growth factor cocktail containing (SCF, FLT3, IL-6, TPO) [26] **(Extended file, Table-3)** 1% Penicillin and Streptomycin for 48 hours. The cells were collected with the media and centrifuged at 130g for 5 mins. The supernatant was collected and centrifuged twice: first at 1000g for 5 min at 4°C, then at 4000g for 15 min at 4°C, and finally at 6000g for 10 min at 4°C. After centrifugation, the supernatant was filtered through a 0.22□µm filter to remove cellular debris, aliquoted, and stored frozen at –80°C for further downstream analysis until use.

### Triple Negative Breast Cancer stem cell sorting and culture

MDA-MB 468, MDA-MB-231, and MCF-7 cell lines, upon reaching 80% confluency, were trypsinized and washed with PBS and then kept in 500 μL of ice-cold FACS dissociation buffer (27). The cells were centrifuged, the supernatant was discarded, and the cell pellets were incubated with CD44 and CD24 antibodies (Invitrogen, Thermo Scientific) antibodies Alexa flour 488 anti-human CD44 and Per CP/Cy5.5 anti□human CD24 (Bio□Legend) (0.5 μg), and their respective isotype control in a staining volume of 100 μl in FACS staining/ sorting buffer. (Supplementary Table; S-5 & S-6) and then subjected to cell sorting using a Bio-Rad S3E cell sorter (Bio-Rad, USA). The cells were sorted as CD44^+^ and CD24^-^ CSCs and were cultured and expanded for further downstream experiments. Details of the antibodies are summarized in supplementary tables S1 and S2.

### Trans-well tropism assay

The trans-well migration assay was performed using trans-well 24-well inserts (8.0 μm, SPL Life Science). HSC tropism towards breast cancer and cancer stem cells (MDAMB-468 CSCs, MDAMB-231 CSCs, MCF-7) and (HaCaT as control cells) was assessed. Breast cancer and cancer stem cells were seeded at 0.05 × 10^6 in 24-well plates overnight (lower chamber). Subsequently, 10,000 HSCs were seeded into the Trans-well chamber (upper chamber), and after 48 hours, the media was removed. The Trans-well was then washed with PBS, fixed with 4% PFA, and permeabilized with chilled ethanol. Crystal Violet (0.5%) staining was performed in the dark for 5 minutes at room temperature, followed by washing and removal of non-migratory cells with a cotton swab. Migratory cells were then quantified using inverted microscopy.

### MTT assay

To assess cell proliferation, MDAMB-468/231 CSCs, MCF-7, and HaCaT cells were plated at a density of 1×10□ cells/200µl and cultured in HSC-CM media for 48 h. The cells were then incubated with MTT for 4 h, dissolved in DMSO, and the absorbance was read at 570 nm using a Beckman Coulter spectrophotometer.

### Annexin□V□FITC/PI apoptosis assay

The apoptosis assay evaluated the ability of HSCs-CM to induce apoptosis in breast cancer cells and cancer stem cells (CSCs) using Annexin-V-FITC and propidium iodide (PI) staining, analyzed via fluorescence microscopy (EVOS M5000) and flow cytometry (Guava EasyCyte). Cells (MDAMB-468/231 CSCs, MCF-7, HaCaT) treated with 50% HSCs-CM for 48h were categorized into healthy, early/late apoptotic, and necrotic populations via quadrant analysis (FCS Express 5/ImageJ) based on Annexin-V/PI staining.

### TUNEL assay

The TUNEL assay assessed HSC-CM-induced apoptosis in MDAMB-468/231 CSCs, MCF-7, and HaCaT cells (1×10□ cells/well, 48h treatment) via DNA fragmentation detection. Cells were fixed, treated with HCl/Triton X-100, stained with TUNEL reaction mix (Invitrogen, ThermoFisher) (fluorescein-dUTP), and analyzed by flow cytometry (Guava EasyCyte), with TUNEL-positive cells quantified as a percentage of total cells.

### Cell cycle analysis

The cell cycle analysis evaluated the effect of HSC-CM on TNBC-CSCs, MCF-7, and HaCaT cells (0.2 × 10^6 cells/well) after 48 h of treatment following serum starvation, using FxCycle™ PI/RNase staining. Flow cytometry (Guava EasyCyte) quantified cell cycle phases (G0/G1, S, G2/M), analyzed via FCS Express 5.0 and GraphPad Prism V7.

### JC-1 staining of mitochondria

The JC-1 assay was performed to access mitochondrial membrane depolarization in TNBC-CSCs, MCF-7, and HaCaT cells (0.3×10□ cells/well) treated with HSC-CM (50:50 ratio) for 48h, using CCCP as a control. JC-1 aggregates (red) and monomers (green) were imaged (EVOS/BIO-RAD), with fluorescence quantified as CTCF (Integrated Density × [Cell Area × Background]) to assess mitochondrial membrane potential.

### Mito-tracker assay

The study assessed mitochondrial membrane potential (ΔΨm) and ROS interplay in TNBC-CSCs, MCF-7, and HaCaT cells (0.3×10□/well) treated with HSC-CM (50:50 ratio, 48h) using MitoTracker Red staining. Fluorescence imaging and flow cytometry were used to quantify mitochondrial integrity and ROS levels, with CCCP (5 μM) serving as a positive control.

### Quantitative real-time PCR (qRT-PCR)

Approximately 1X10^6^ TNBC-CSCs, MCF-7, and HaCaT were harvested with TrypLE (**Invitrogen, ThermoFisher**), washed with DPBS (Invitrogen, ThermoFisher), and subjected to total RNA isolation using the Trizol method (Taqara, USA). RNA quantification was performed, followed by cDNA synthesis using the iSCRIPT™ cDNA synthesis kit with 1 μg of RNA. This was then followed by qRT-PCR using SSO-Fast™ Eva Green Supermix. The mRNA expressions (Ct values) were normalized to GAPDH, generating δCt values. δCt values exceeding 18 were considered as not expressed (NE). The details of the primers are provided in the Supplementary Extended file (Table 1).

**Table 1:** Mitochondrial gene expression profile of TNBC-CSCs, MCF-7, and HaCaT control cells after post-HSCs-CM derived treatment. Statistical analysis, 2 Way-ANOVA with Sidak’s multiple comparison test, p-value *p < 0.05 considered statistically significant. Upregulated is denoted as Up, and Downregulated is denoted as Down.

**Fig: 7**
| Genes | Cell lines |  |  |  |
| --- | --- | --- | --- | --- |
|  | HaCaT | MDAMB-468<br>(CSCs) | MDAMB-231(CSCs) | MCF-7 |
| **SCO2** | 1.229484; Up | 0.026; Down | 0.113247; Down | 0.687364; Down |
| **SURF1** | 3.465249; Up | 0.021; Down | 0.05634; Down | 1.164545 |
| **COA6** | 2.468206; Up | 0.024; Down | 0.083191; Down | 0.442759; Down |
| **MTCO1** | 2.44957; Up | 0.69; Down | 0.255923; Down | 0.523987; Down |
| **MTCO2** | 1.435491; Up | 0.27; Down | 0.086989; Down | 0.047093; Down |
| **COX4I1** | 2.385001; Up | 0.087; Down | 0.006802; Down | 0.208186; Down |
| **DRP1** | 1.644924; Up | 8.3; Up | 5.617131; Up | 4.412726; Up |
| **FIS1** | 1.81742; Up | 5.93; Up | 14.44591; Up | 2.040386; Up |
| **OPA1** | 2.596065; Up | 0.71; Down | 0.001033; Down | 0.034241; Down |
| **Mfn-1** | 2.40606; Up | 0.19; Down | 0.094578; Down | 0.53203; Down |
| **Mfn-2** | 2.668191; Up | 0.24; Down | 0.319043; Down | 0.218154; Down |
| **Parkin-2** | 2.05569; Up | 4.17; Up | 1.543751; Up | 4.649666; Up |
| **TOMM-20** | 7.404047; Up | 29.41 | 65.28244; Up | 11.21747; Up |
| **TOMM-22** | 4.694335; Up | 19.76; Up | 3.91783; Up | 20.58699; Up |
| **TOMM-70** | 1.947912; Up | 4.42; Up | 9.74616; Up | 4.522016; Up |

### Spheroid formation of TNBC CSCs in HSCs-CM

To check the spheroid formation ability of CSCs in the presence of HSCs-CM. Cells were harvested and used to induce spheroid formation in MDAMB-468 CSCs, MDAMB-231 CSCs, MCF-7, and HaCaT cells. The cells were cultured in cell-specific media and HSC-derived conditioned media (in a 50:50 ratio) using the hanging drop method, with approximately 800 cells per drop, and incubated for 48 h. After 48 hours, spheroids were observed under the EVOS M5000 imager (Thermo Fisher). Images were captured and analyzed.

### Proteomics studies of migrated Hematopoietic Stem Cells

Migrated single-cell proteomic analysis was performed to investigate the role of HSCs in the tumor microenvironment. Briefly, after co-culture of HSCs, migrated cells were collected, and proteins were extracted in SDS buffer (BCA-quantified). The proteins were then digested with trypsin and analyzed via LC-MS/MS (Orbitrap Fusion Tribrid, 120K resolution, 350-1500 m/z). Data processed with DIA-NN against UniProt (1% FDR) identified differentially expressed proteins after median normalization, considering carbamidomethylation, oxidation, and acetylation modifications.

### Metabolomics studies of CM-treated TNBC and MCF-7 cells

Metabolomic profiling of TNBC-CSCs, MCF-7, and HaCaT cells post-HSC-CM treatment (48h) involved metabolite extraction using 80% chilled methanol, centrifugation, SpeedVac drying, and LC-MS/MS analysis (Q Exactive Orbitrap, 140K resolution, 100-700 m/z). Data processed via Compound Discoverer 3.2 identified metabolites with HCD fragmentation (NCE 35%), dynamic exclusion, and 35K resolution for fragment ions (28).

### Western blotting

For Western blotting, TNBC-CSCs, MCF-7, and HaCaT cells were lysed in Tris/SDS buffer, with proteins (20µg) separated on 10% SDS-PAGE, transferred to nitrocellulose, and probed with primary/secondary antibodies. Bands detected via ECL (ChemiDoc) were quantified (ImageJ) and normalized to β-actin as a loading control.

### Statistical Analysis

All experiments were performed as three biological replicates and statistical analysis was conducted using GraphPad Prism 8 (GraphPad Software, San Diego, USA) and R (version 4.2.2). Metabolomics data were acquired in positive mode with a single replicate. The results were expressed as Mean ± SEM. Results with a p-value of less than 0.05 were considered significant. One-way ANOVA was performed to compare more than two groups unless mentioned otherwise. Two-way ANOVA with Sidak’s multiple comparison test was performed for the multiple groups. The R program, SR plot, and Metaboanalyst (V 6.0) were used to generate the heat map, chord diagram, and volcano plot for the OMICS data analysis.

## Results

### HSCs maintained stem cell properties after sorting

HL-60 cells cultured in RPMI media supplemented with SCF, FLT3, IL-6, and TPO were analyzed by flow cytometry. Results showed 56.9% expressed CD34 and 84.07% CD45, with co-expression of both markers. Minimal expression (≤1.79%) of CD105, CD19, CD14, and HLA-DR was observed compared to **(S1; a and c)** controls. CD34^+^/CD45^+^ hematopoietic stem cells (HSCs) were sorted using fluorescence-activated cell sorting (**S2; a**), expanded in growth factor-enriched media, and used for subsequent studies.

Human breast cancer stem cells, sorted for CD44^+^/24^-^ from the MDA-MB-468 and MDA-MB-231 cell lines used in our previous study, were revived from liquid nitrogen and utilized in the current experiments (27). All three cell lines were subjected to flow cytometry analysis further to confirm the percentage of the stem cell population. The results indicated that MDAMB-468 cells maintained the stem cells properties and expressed CD44 (98.89%) and CD24 (0.55%), MDAMB-231 cells expressed CD44 (99.60%) and CD24 (3.91%), and MCF-7 cells expressed, very low CD44 (0.73%) and CD24 (0.26%) (**S1; b and d**). All the cells mentioned above, sorted and cultured, were used for our study’s downstream experiments.

### HSCs exhibited tropism towards the TNBC-CSCs and MCF-7 HER2^+^ cancer cells

To determine the tropism properties of HSCs towards TNBC-CSCs, a co-culture-based tropism assay was performed, and the results indicated that migration was observed (as indicated by the crystal violet stain on the membrane, Figure 1a), with the migration of HSCs varying between the experimental and control groups. The total number of HSCs migrated through the trans-well insert towards MDAMB-468 CSCs was 30 cells/unit area (a.u.), MDAMB-231 CSCs were recorded as 28 cells/unit area, and MCF-7 was 22 cells/area (**Figure 1a & 1b**). Moreover, the HSCs showed minimal migration towards control HaCaT cells, with only 4 cells per unit area, indicating an exclusive preference for cancer and cancer stem cells, thereby validating the tumor tropism of HSCs (**Figure 1a-b**). Furthermore, flow cytometry-based corroboration of the migrated HSCs indicated a presence of 98.59% HSCs migrating towards MDAMB-468 CSCs, 92.47% HSCs migrating towards MDAMB-231 CSCs and 95.72% HSCs migrating towards MCF-7 CSCs. In contrast, only 5% of HSCs had migrated towards the control cell line HaCaT, thereby indicating only 5% HSC tropism in the HaCaT cell line used as a control in this study **(Figure 1c-d)** in the presence of HSCs.

**Figure 1:**
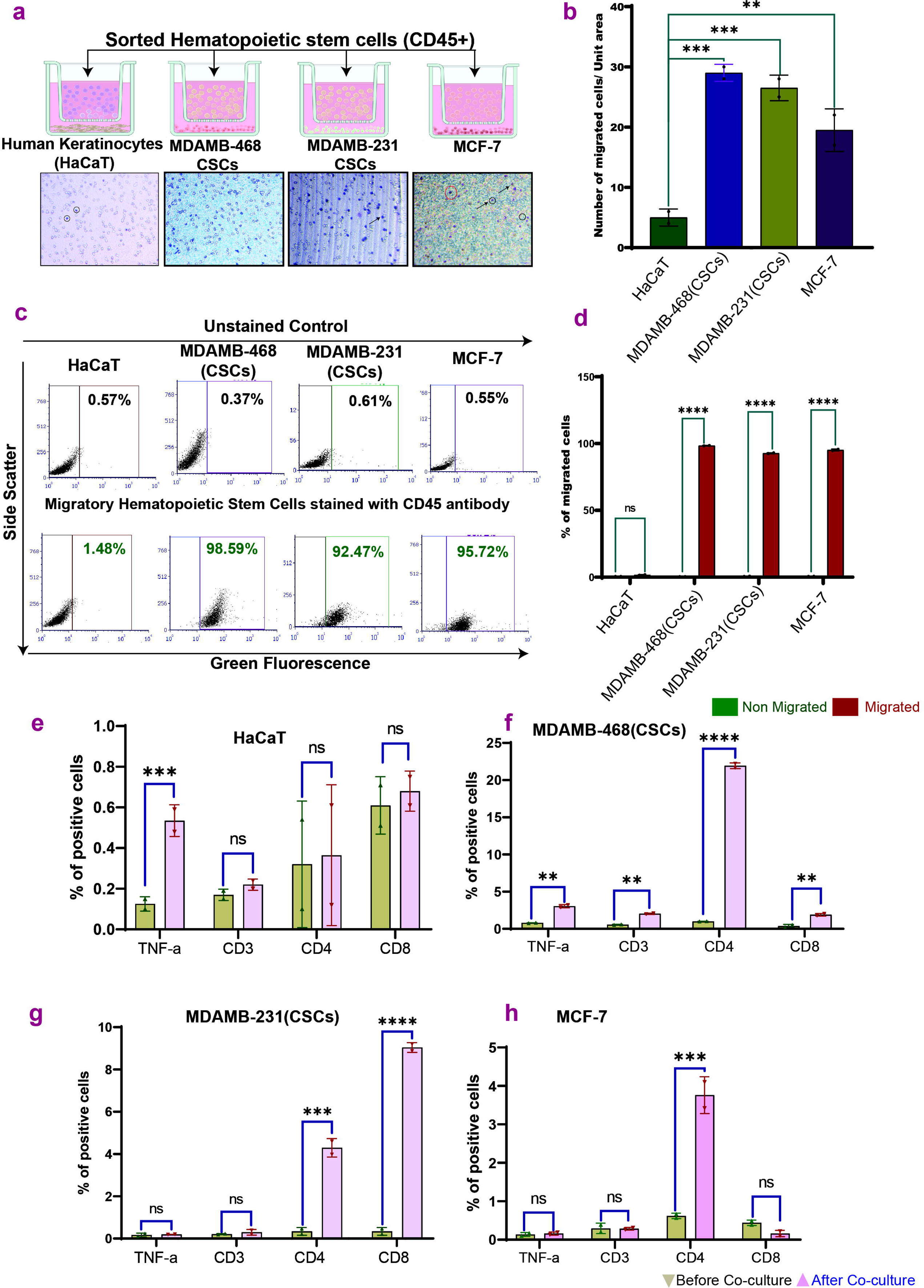
Tropism of HSCs on TNBC-CSCs, MCF-7, and HaCaT control cells. Figure 1, a) is the graphical representation of the tropism assay. Figure 1.a) represents the crystal violet staining of migrated HSCs after co-culture with all TNBC cell lines and MCF-7 cells (arrow marks showing the migrated cells), and **b)** the bar graph represents the quantification of migratory HSCs towards HaCaT, TNBC-CSCs and MCF-7 cells. **Figure1 c)** represents the flow cytometry validation of HSCs migratory cells towards HaCaT and TNBC CSCs and MCF-7 cells, and **d)** bar graph represents the quantification of HSCs migratory cells towards the TNBC CSCs, MCF-7 and control cell line HaCaT. **Figure1, e-h)** shows the flow cytometry-based T lymphocyte and inflammation markers profiling of migrated HSCs and differentiation towards CD-3, CD-4, and CD-8 cells in TNBC CSCs, MCF-7, and HaCaT control cells. Values are represented as Mean ± SEM. *p < 0.05, **p < 0.01, ***p < 0.001, ****p < 0.0001.

### Migrated HSCs showed upregulated proteins involved in cell adhesion and migration

To further explore HSC tropism properties, migrated HSCs were collected from the bottom chamber of TNBC-CSCs and MCF-7 and single-cell type proteomics studies were performed. Our analysis uncovered a complex proteomic landscape, identifying 2757 proteins, including key regulatory proteins. Notably, 381 proteins were significantly dysregulated, with 194 upregulated and 187 downregulated. Upon analysis, we found that the maximum dysregulated proteins are involved in cell adhesion, migration, and cytoskeleton remodeling processes **(Fig 2),** again indicating the abilities of HSCs to modulate such critical cellular processes, possibly governing cell metastasis in the tumor microenvironment. Moreover, the volcano plot showed that the dysregulated proteins are involved in response to stimulus, protein metabolic process, and protein translational and structural constituents of cytoskeleton processes **(S-2; b-e)**.

**Figure 2:**
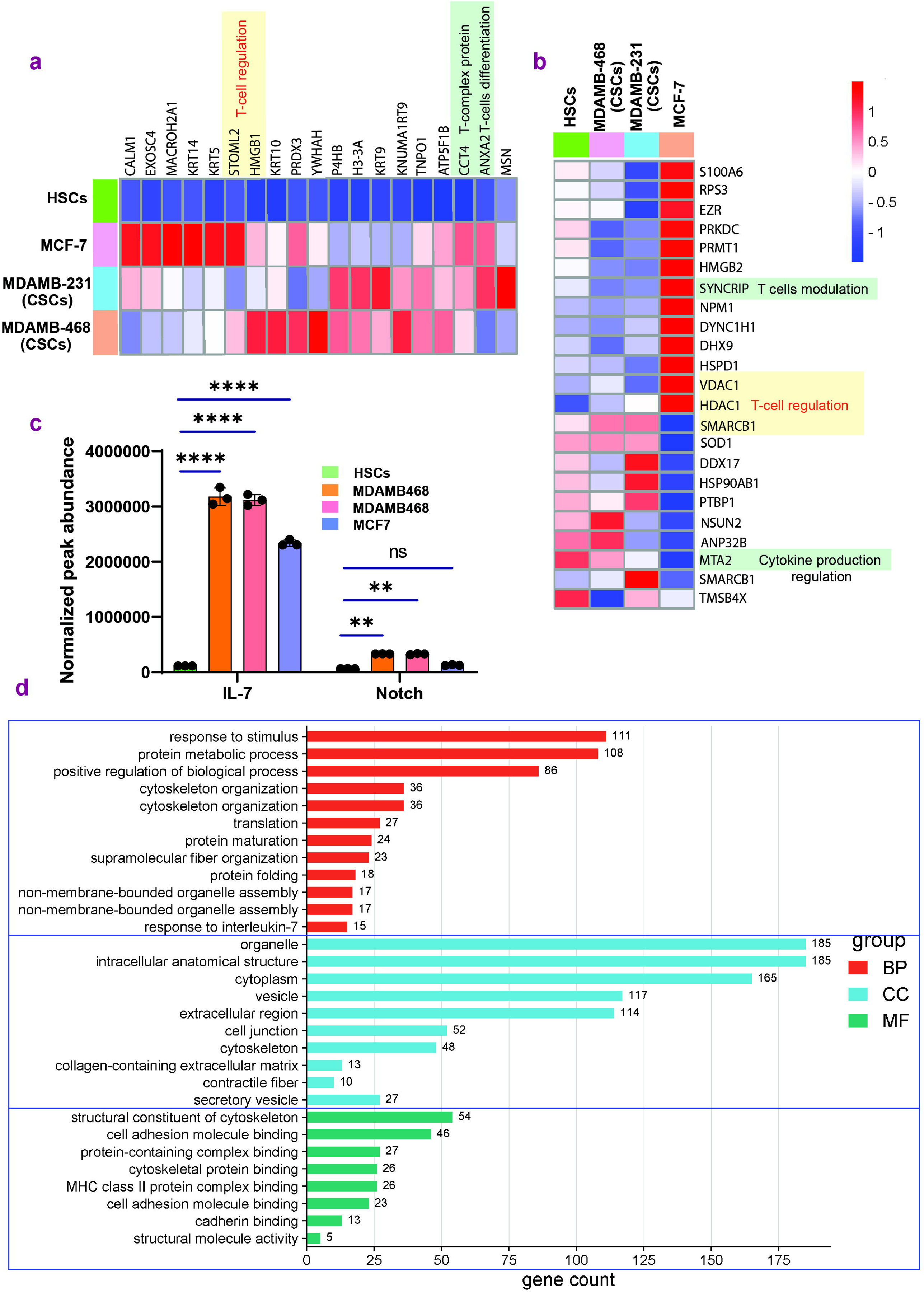
Proteomics profiling of the migrated HSCs towards TNBC-CSCs and MCF-7 cells. **Figure 2, a-b)** Heatmap of dysregulated proteins of migrated HSCs towards TNBC-CSCs and MCF-7 cells revealed T-cells differentiation and T-cells regulation associated signaling pathway. **c)** normalized peak expression of IL-7 and Notch signaling protein in MDAMB-468 (CSCs), MDAMB-231 (CSCs) and MCF-7 cells. (p□<□0.05). **d)** Gene ontology of the migrated HSCs and altered proteins and their involvement in various biological pathways, molecular functions and cellular components.

### Differentiation of HSCs into cytotoxic T cells when co-cultured with TNBC-CSCs

After HSCs migrated toward TNBC-CSCs, their differentiation into cytotoxic T cells was analyzed in flow cytometry. Co-culture with TNBC-CSCs triggered HSC differentiation into CD4^+^ (20% for MDAMB-468, 6% for MDAMB-231, 4% for MCF-7) and CD8^+^ T cells (3%, 9%, 2%, respectively), with varying statistical significance (MDAMB-468: CD4^+^ p<0.0001; MDAMB-231: CD8^+^ p<0.0001) (**Figure 1e**). No differentiation occurred with HaCaT controls, confirming tumor microenvironment specificity **(Fig. 1e-h, S3)**.

Migrated single cells proteomics revealed upregulated T cell-related proteins (e.g., IL-7, Notch) in HSCs exposed to TNBC-CSCs/MCF-7 versus controls, aligning with their differentiation capacity **(Fig. 2a-d)** (29–31). These findings highlight HSC plasticity in adapting to the breast cancer microenvironment, driven by molecular cues such as the IL-7/Notch protein **(Figure 2c).**

### HSCs-derived conditioned media drastically impaired the cell viability of TNBC-CSCs and MCF-7 cells

Our previous experiments have proven that HSCs differentiate into cytotoxic T lymphocytes in the breast cancer milieu; we hypothesized that secretory factors from HSCs affect TNBC-CSCs/MCF7. Metabolomics of HSC-derived conditioned media revealed that those enriched in metabolites linked to amino acid, lactose, sphingolipid, and fatty acid pathways significantly reduced viability (32). As the HSCs-CM was found to be abundant in metabolites, we further deployed the hematopoietic stem cells derived conditioned media for the cellular viability studies and results indicated drastically reduced the cellular viability of TNBC-CSCs (MDAMB-468: 48h p<0.01; MDAMB-231: 48h p<0.0001) and MCF-7 cells (48h p<0.0001), with minimal effect on HaCaT controls **(ns, S4a-d)**. Further, live-dead staining confirmed HSC-CM induced 79-90% cell death in TNBC-CSCs and 45.94% in MCF-7, while HaCaT viability remained unchanged **(S5)**, demonstrating selective cytotoxicity toward breast cancer cells.

### HSCs CM altered the metabolites in the TNBC-CSCs and MCF-7 cells

Next, we performed global metabolomics studies on CM-treated TNBC-CSCs and MCF cells to understand the alterations in metabolite levels. Pathway enrichment and gene ontology results revealed that various amino acid-mediated pathways are dysregulated **(S-6, a-b)**. Dysregulation of arginine biosynthesis and metabolism may lead to the apoptosis process and other branched-chain amino acids such as (valine, leucine, and isoleucine) BCCAs, which play an important role in energy metabolism and its alteration, leading to mitochondrial dysregulation and further apoptosis.

Furthermore, metabolic pathway analysis of differentially expressed metabolites was performed using MetaboAnalyst (v6.0) for the CM-treated TNBC CSCs and MCF-7 cells. Several pathways were significantly enriched (FDR ≤ 0.05), including arginine and proline metabolism, purine metabolism, pyrimidine metabolism, and phenylalanine metabolism, among others **(S-7; a-b)**. Further, metabolite classification revealed that various metabolites targeted various proteins, transcription factors, and biological pathways **(S-6)**

### Inactivation of cell cycle checkpoints when CSCs and MCF-7 cultured in HSC-CM

Next, we wanted to investigate the responses of TNBC-CSCs, MCF-7, and HaCaT cells when cultured in HSCs-CM. HSC-CM disrupted cell cycle dynamics in TNBC-CSCs and MCF-7, causing S-phase reduction and G2/M arrest in MDAMB-468 (S-phase: 69.89%→24.07%; G2/M: 11.22%→49.10%) and MDAMB-231 (S-phase: 42.65%→25.23%; G2/M: 6.79%→10.78%). MCF-7 cells showed G0/G1 arrest (G0/G1: 37.93%→54.41%), while HaCaT controls exhibited increased S-phase (47.07%→58.97%) (Fig. 3a, S7b).

**Figure 3:**
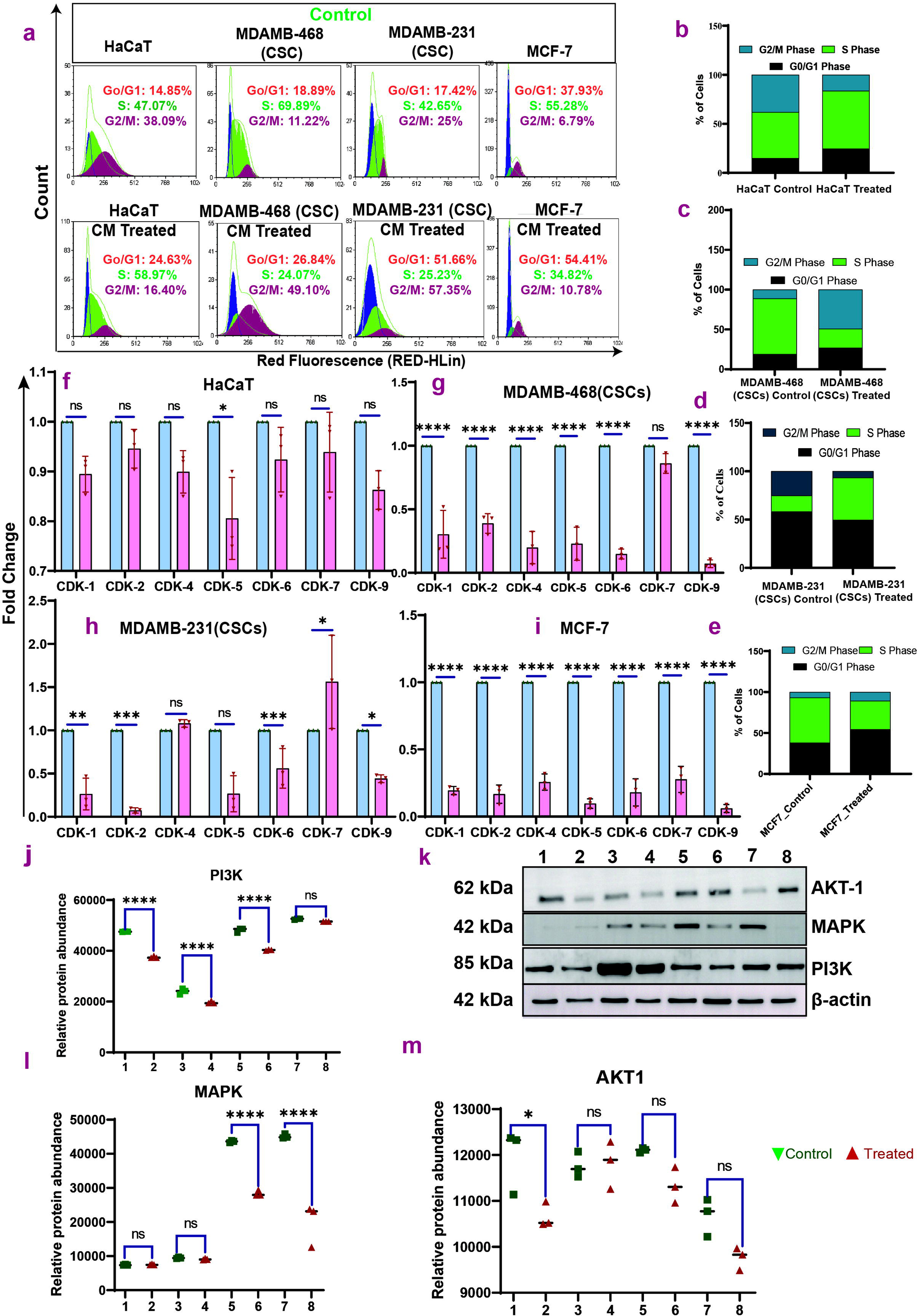
**a) Cell cycle analysis of TNBC cell lines (**MDAMB-468, MDAMB231), MCF-7 and HaCaT cells after being treated with HSCs-derived conditioned media for 48 hours. Results indicated that HSCs-derived conditioned media induces synthesis phase arrest in MDAMB-468 (CSCs) and MCF-7 cells and G2/M phase arrest in MDAMB-231(CSCs) cells, but no changes occurred in HaCaT cells. **Figure 3; b-e):** Gene expression profiling of MDAMB-468 (CD44^+^), MDAMB231(CSCs), MCF-7, and HaCaT cells after 48 hours cultured with HSCs derived conditioned media. HSC-derived conditioned media downregulates the CDKs gene and suppresses the proliferation of TNBC CSCs, but in HaCaT control cells, no changes occur. **Fig-3; f-i)** Western blot-based protein expression profile of CM-treated TNBC-CSCs, MCF-7, and HaCaT cells (AKT-1, MAPK, and PI3K). Values are represented as Mean ± SEM. *p < 0.05, **p < 0.01, ***p < 0.001, ****p < 0.0001.

CDKs (1,2,4,5,6,7,9) were significantly downregulated in TNBC-CSCs and MCF-7 (most p<0.001) but largely unaffected in HaCaT **(Fig. 3b-e)**. **Moreover, metabolomics enrichment data hence pointed out the possibility that the** metabolomics linked HSC-CM to the inhibition of oncogene transcription by suppressing polymerase-II phosphorylation. Western blot confirmed downregulation of PI3K, AKT-1, and MAPK in cancer cells, aligning with cell cycle and CDK data **(Fig. 3f-i, S7a)** (33). These pathways regulate cyclins/CDKs and proliferation, underscoring HSC-CM’s role in targeting cancer cell cycle mechanisms.

### HSC-CM-induced double-strand DNA breaks in TNBC-CSCs

Since HSC-CM inhibited the cell cycle in TNBC-CSCs and MCF-7 cells, we next checked DNA breakage in these cells. HSC-CM induced significant DNA damage in TNBC-CSCs and MCF-7, with TUNEL-positive cells at 65.39% (MDAMB-468), 58.44% (MDAMB-231), and 76.84% (MCF-7) (p<0.0001), versus none in HaCaT controls **(Fig. 4a-b, S8)**.

**Figure 4:**
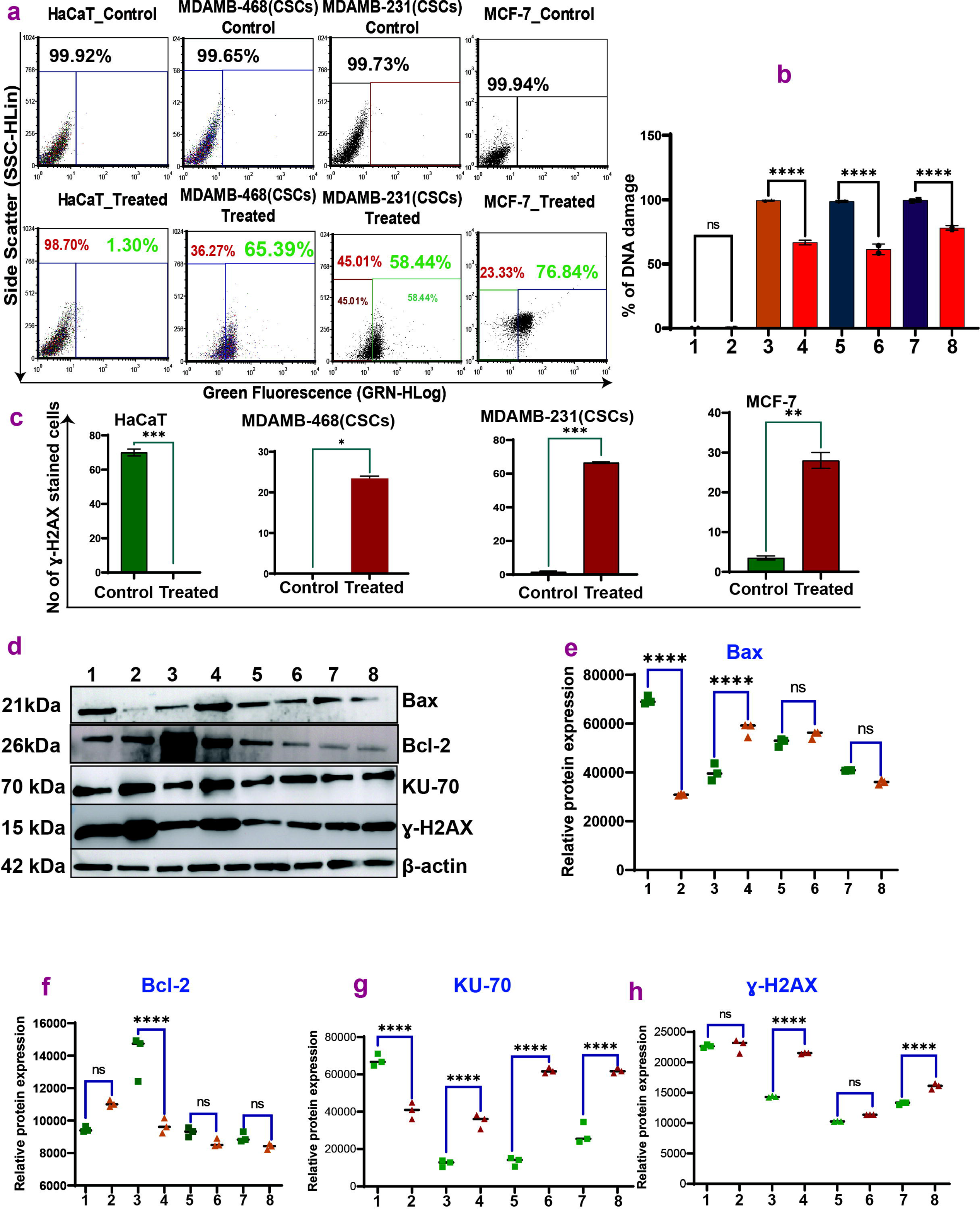
**a) Flow cytometry-based TUNEL assay for double-stranded DNA breaks (DSB) in TNBC-CSCs (MDAMB-468, MDAMB-231) & MCF7 cells** post 48 hours of CM treatment and **b)** quantification of the DSB assay in TNBC-CSCS and MCF7 cells. **Figure-4; c)** bar diagram represents the γ-H2AX stained cells of the immunocytochemistry fluorescent images (**Supplementary Figure S-10; blue representing DAPI, green is γ-H2AX stained cells**) of the TNBC CSCs and HaCaT after 48 h post-CM treatment. Scalebar: 30µm. Figure 4**; d)** Western blot of key apoptotic proteins and DNA damage proteins in the CM treated TNBC-CSCs, MCF-7, and HaCaT cells and **e-k)** bar graph representing the relative protein expression of the apoptotic related proteins in the CM treated cells. 1=HaCaT control, 2= HaCaT Treated, 3=MDAMB-468(CSCs) Control, 4=MDAMB-468(CSCs) Treated, 5=MDAMB231(CSCs) Control, 6=MDAMB-231(CSCs) Treated, 7=MCF-7 Control, 8=MCF-7 Treated.

Furthermore, we performed immunostaining and Western blot to evaluate γ-H2AX expression and relative protein expression in HSCs-CM-treated TNBC-CSCs, MCF-7, and HaCaT cells. Results revealed that there was a significant increase in the expression of γ-H2AX post 48h treatment in the MDAMB-468 (22 ±cells), MDAMB-231 (65± cells) and MCF-7 (27 ±cells) when compared with control HaCaT cells which did not show any γ-H2AX positive cells (**Fig. 4c, h & S9; a-c**). KU70, another protein primarily involved in non-homologous end joining (NHEJ), also increased upon CM treatment compared with non-treated and HaCaT control cells **(Fig-4; g)**. The upregulation of KU70 protein expression further validated the DNA damage that occurred in the TNBC-CSCs upon CM treatment.

### HSC-CM causes apoptosis in the TNBC-CSCs

Next, we investigated the apoptosis pathway in TNBC-CSCs and MCF-7 cells induced by HSCs-CM for 24 h and 48 h. HSC-CM induced apoptosis in TNBC-CSCs and MCF-7 cells after 48h, causing necrosis in MDAMB-468 (70.94%) and MDAMB-231 (96.31%) and late apoptosis in MCF-7 (34.77%). HaCaT controls showed no discernible changes when cultured in HSCs-CM (Fig. 5a, S10). Subsequently, we examined the mRNA and protein expression of key apoptosis-related genes, including p53, Bcl-2, and Bax, in these cells. Gene expression analysis revealed upregulation of pro-apoptotic markers: p53 (up to 18-fold in TNBC-CSCs, 12-fold in MCF-7), Bax (up to 58-fold in TNBC-CSCs, 25-fold in MCF-7), and FOXO3a (immunofluorescence), alongside downregulation of anti-apoptotic Bcl-2 (≤0.6-fold) **(Fig. 5b-f, i-m, S9c)**. Gene expression results were similar to the protein expression of both Bax **(Figure 4; d-e)** and Bcl-2 **(Figure 4; d & f)** protein in CM-treated TNBC, HER2^+^ control cells, thereby again indicating that HSCs-derived CM directly affects apoptosis gene and p53, subsequently modulating the expression of pro and anti-apoptotic genes and hence apoptosis, exclusively in TNBC CSCs and MCF-7 cells.

**Figure 5:**
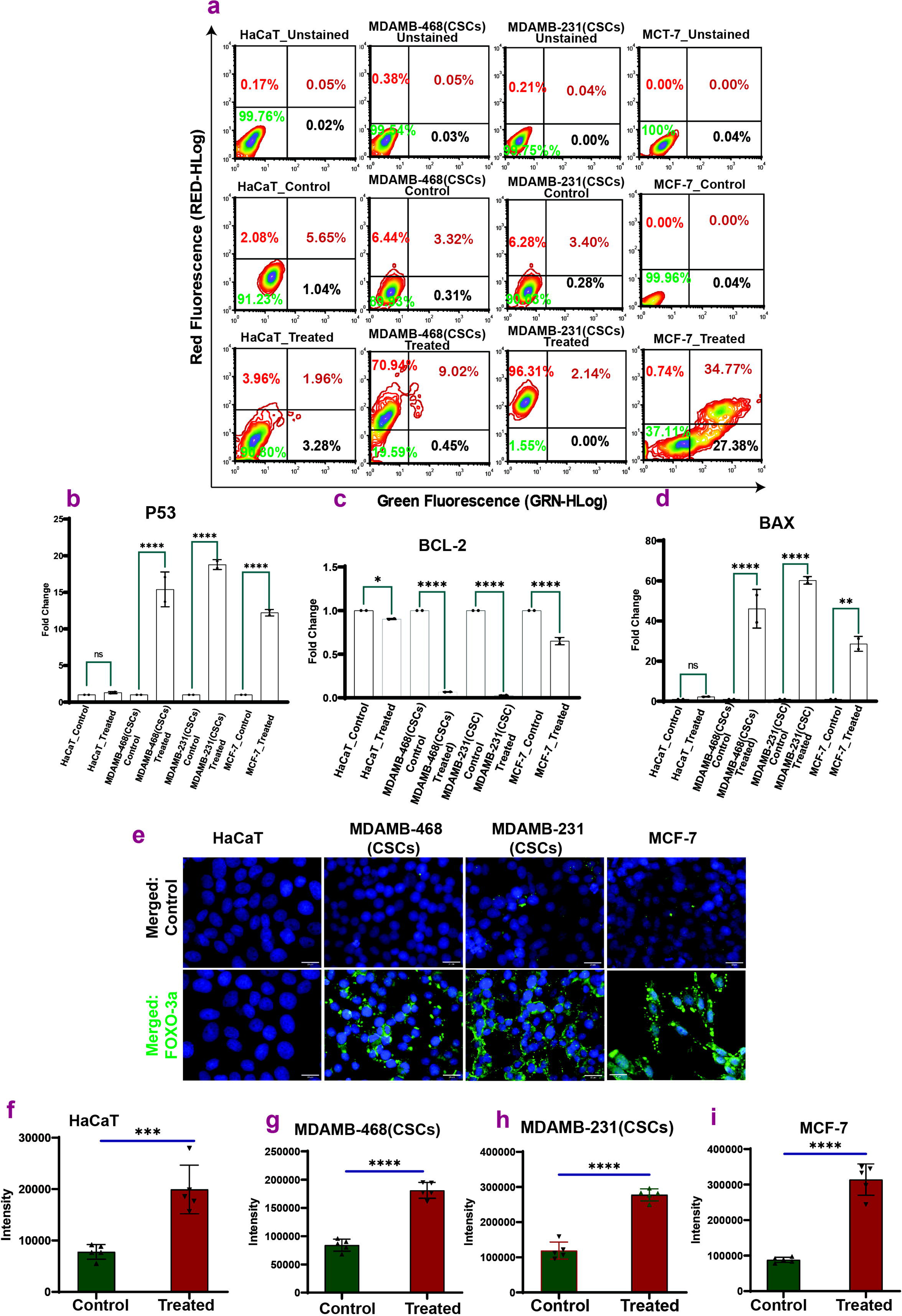
Apoptosis and gene expression in responses to HSCs-derived CM in TNBC Stem Cells and MCF-7 cells. **a)** Flow cytometry analysis of TNBC stem cells and MCF-7 cells treated with HSCs-derived conditioned media (CM) for 48 hours with no significant changes in apoptosis induction in control HaCaT cells, MDAMB-468 (CSCs) exhibit late apoptosis (11.94%) and necrosis (5%). MDAMB-231(CSCs) predominantly undergo early apoptosis (94.16%). MCF-7 cells show increased late apoptosis (18.96%) and necrosis (17.17%). (Fig-5: b-d) gene expression profile of the P53 (fig-5: b), BAX (fig-5: d) and Bcl-2 (fig-5: c) after treatment of HSCs derived CM on TNBC CSCs, MCF-7 and HaCaT cells. **e)** Immunocytochemistry imaging based FoXo3a expression in CM treated TNBC-CSCs, MCF-7 and HaCaT cells and **f-i)** bar graph represents the intensity of the FoXo3a expression in the mentioned cells. Values are represented as Mean ± SEM. *p < 0.05, **p < 0.01, ***p < 0.001, ****p < 0.0001.

These changes, absent in HaCaT, confirm HSC-CM selectively triggers apoptosis in cancer cells via p53/Bax activation and Bcl-2 suppression. It is, hence, essential to emphasize that FOXO3a is involved in the pro-apoptotic pathway(29,34), and the upregulation of this transcription factor further established the CM-induced apoptosis in the TNBC-CSCs and MCF-7 cells **(Figure 5; i-m and S-9c)**.

### HSC-CM decreases mitochondrial morphology and quality in TNBC-CSCs

As evidenced in the previous sections regarding the apoptosis-inducing potential of HSC-CM in the TNBC CSCs and MCF-7 cells, we further investigated the impact of HSC-CM on the mitochondrial health of the aforementioned cells, including the HaCaT control cells. HSC-CM disrupted mitochondrial health in TNBC-CSCs and MCF-7 cells, causing fragmentation, perinuclear aggregation, and dysregulated fission/fusion (Fig. 6b-c). In contrast, HaCaT controls retained intact mitochondrial networks post-treatment (Fig. 6c-d, S11). These structural disruptions suggest that HSC-CM impairs mitochondrial function (e.g., ATP loss), potentially driving apoptosis in cancer cells.

**Figure 6:**
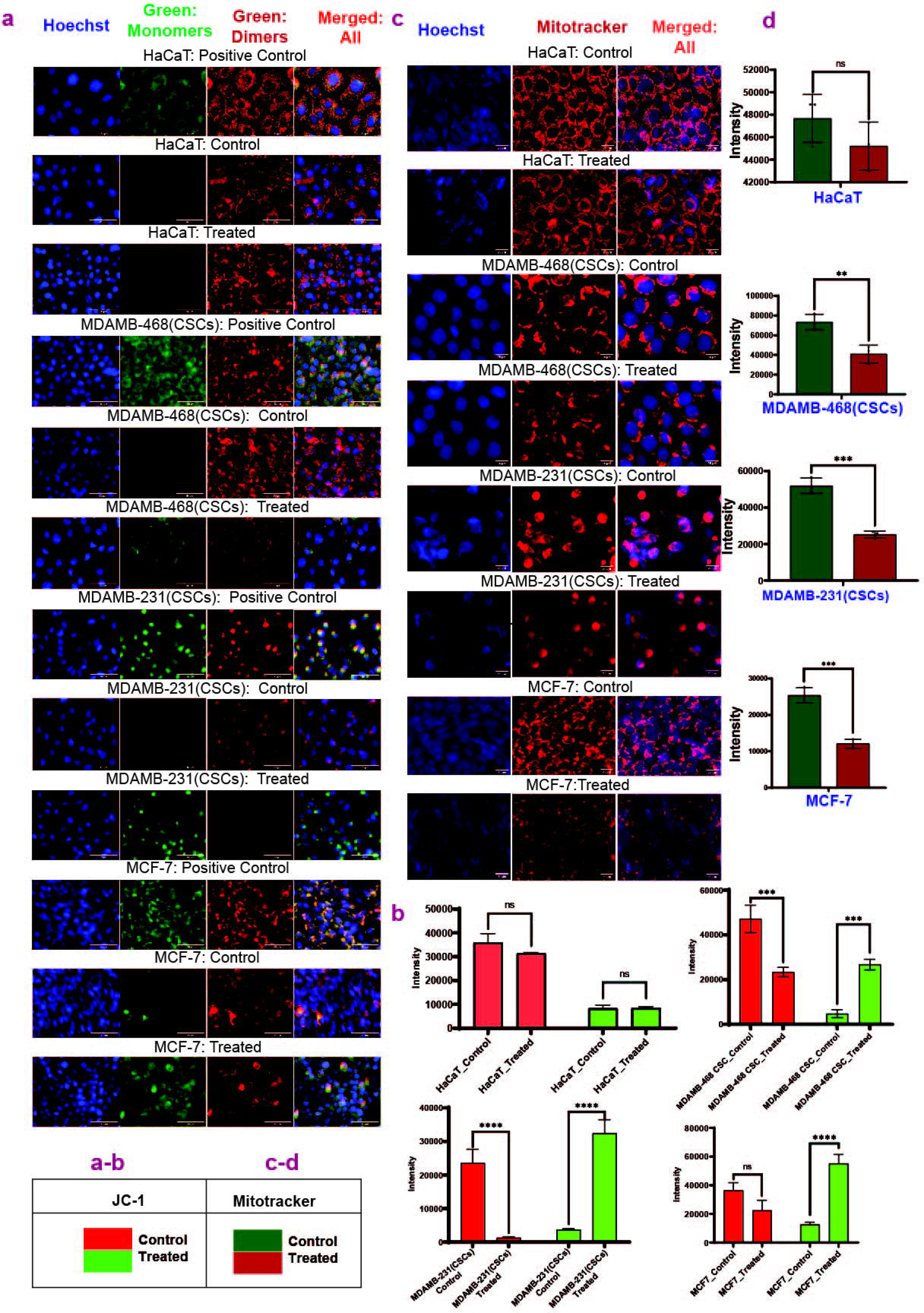
HSCs CM alters the mitochondrial dynamics in the TNBC-CSCs and MCF-7 cells. Figure 6; a) Fluorescence images of JC1 staining in CM treated TNBC-CSCs, MCF-7 and HaCaT control cells; Scale bar: 50μm, **b)** Bar graph representing the quantification of intensity of JC1-green and JC1-red in CM treated cells. **c)** Fluorescence images of mitrotracker staining of TNBC-CSCs, MCF-7 and HaCaT control cells, **d)** Quantification of the intensity of active mitochondria in mentioned cells bar: 20μm.Values are represented as Mean ± SEM. *p < 0.05, **p < 0.01, ***p < 0.001, ****p < 0.0001.

### HSC-CM significantly reduced the mitochondrial membrane potential in TNBC cancer stem cells

Next, we evaluated the mitochondrial membrane potential (JC-1 staining) of all the aforementioned cells and the control HaCaT cells upon HSC-CM treatment. Results indicated treatment with HSC-CM on to the TNBC-CSCs and MCF-7 cells caused an increase in the accumulation of JC1-green monomers as an indication of an increase in the depolarization of the mitochondrial membrane or decrease in mitochondrial membrane potential (**Fig. 6 a-b**). CCCP was used as a positive control to indicate the accumulation of unhealthy mitochondria, also exhibiting green monomer abundance, as observed using JC1 staining. Further, when control HaCaT cells were treated with HSC-CM, there was no accumulation of JC1-green monomers, indicating that HSC-CM did not affect the depolarization of the mitochondrial membrane in the control HaCaT cells (**Fig. 6a**).

### Mitochondrial gene and protein expression changes in TNBC-CSCs when cultured in HSCs-CM

Gene expression results showed that mitochondrial fission genes (Drp-1, Fis-1, Parkin-2) were upregulated and fusion genes (Mfn-1/2, OPA-1) downregulated in cancer cells, alongside decreased COA6 (linked to cytochrome c assembly). HaCaT showed no significant gene changes, which confirms HSC-CM selectively disrupts mitochondrial dynamics in cancer cells, driving fragmentation/dysfunction (Fig. 7a, S12, S14, Table 1).

**Figure 7:**
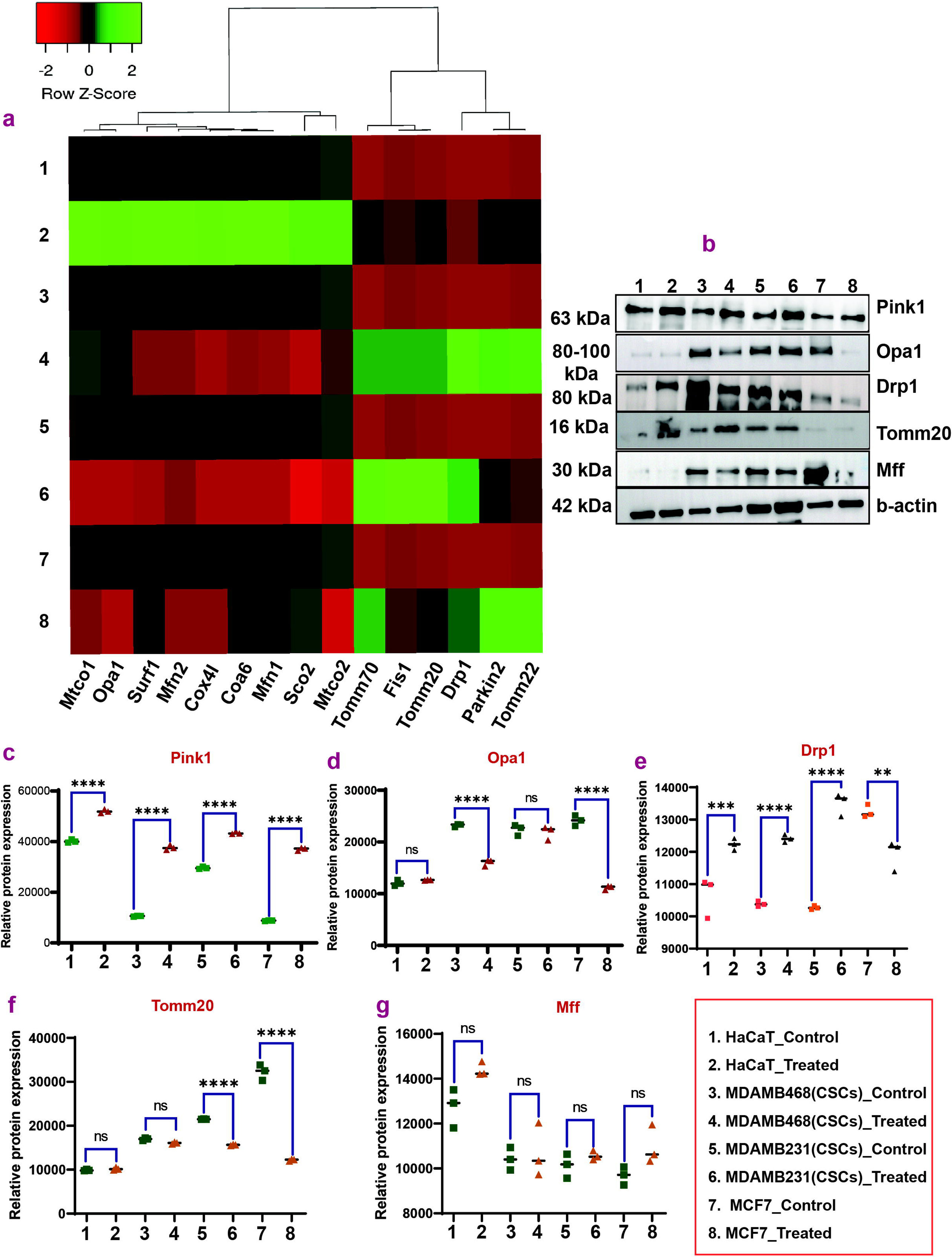
Mitochondrial gene expression profiling after HSCs derived CM treatment on TNBC-CSCs. Figure 7**: A) The heatmap shows the mRNA expression of** mitochondrial dynamics, mitochondrial membrane proteins, and cytochrome oxidoreductase in CM-treated HaCaT, TNBC-CSCs, and MCF-7 cells. **b)** Western blot of the major mitochondrial dynamics proteins and their expression compared after CM treatment, **c-g)** quantification of the protein expression mitochondrial dynamics proteins (Pink-1, OPA-1, Drp-1, Tomm-20, MFF). 1=HaCaT control, 2= HaCaT Treated, 3=MDAMB-468(CSCs) Control, 4=MDAMB-468(CSCs) Treated, 5=MDAMB231(CSCs) Control, 6=MDAMB-231(CSCs) Treated, 7=MCF-7 Control, 8=MCF-7 Treated. Values are represented as Mean ± SEM. *p < 0.05, **p < 0.01, ***p < 0.001, ****p < 0.0001

### Upregulation of mitophagy mitochondrial membrane Proteins upon HSC-CM treatment

Furthermore, all assessed translocase of the outer mitochondrial membrane (TOMM) proteins exhibited significant upregulation in TNBC-CSCs and MCF-7 cells (Table 1). **(Figure 7a & S-13 and Table-1)**. Next, we investigated the major genes involved in maintaining the assembly of cytochrome C oxidase and its subunits, and the results revealed that HSC-CM substantially dysregulates their role and function. Results indicated that the SCO2, SURF-1, COA6, MTCO1, MTCO2, and COX4I are significantly downregulated compared to the control HaCaT (MDAMB-468; p < 0.0001; MDAMB-231, p < 0.0001, MCF-7 p>0.05 and HaCaT, p>0.05) **(Figure 7a & S-14 and table-1).**

### HSC-CM treatment disrupted the spheroid-forming ability in TNBC-CSCs and MCF-7 cells

HSC-CM disrupted spheroid formation in TNBC-CSCs and MCF-7 cells **(Fig. 8a-e)**, which was linked to the downregulation of adhesion/ECM-related genes, including SP-1 (critical for CXCR4-mediated spheroid stability), IBSP, Fibronectin, N-cadherin, SPARC, and ALDH1A1. These genes regulate cell-cell interactions, ECM composition, and CSC enrichment, which are essential for spheroid integrity. TNBC-CSCs and MCF-7 showed significant decreases (p<0.0001), while HaCaT controls remained unaffected (p>0.05). SP-1 reduction (0.02–0.004-fold) impaired cell adhesion, and ALDH1A1 suppression diminished CSC-like states. IBSP (integrin binding) and Fibronectin/N-cadherin (ECM adhesion) downregulation further destabilized the spheroid structure (35–39)37-41. SPARC’s ECM modulation and ALDH1A1’s CSC role highlight their collective impact. Results confirm HSC-CM selectively targets cancer cells by crippling spheroid-associated pathways, sparing non-cancerous HaCaT cells (40–43) **(Fig. 8f-i)**.

**Figure 8:**
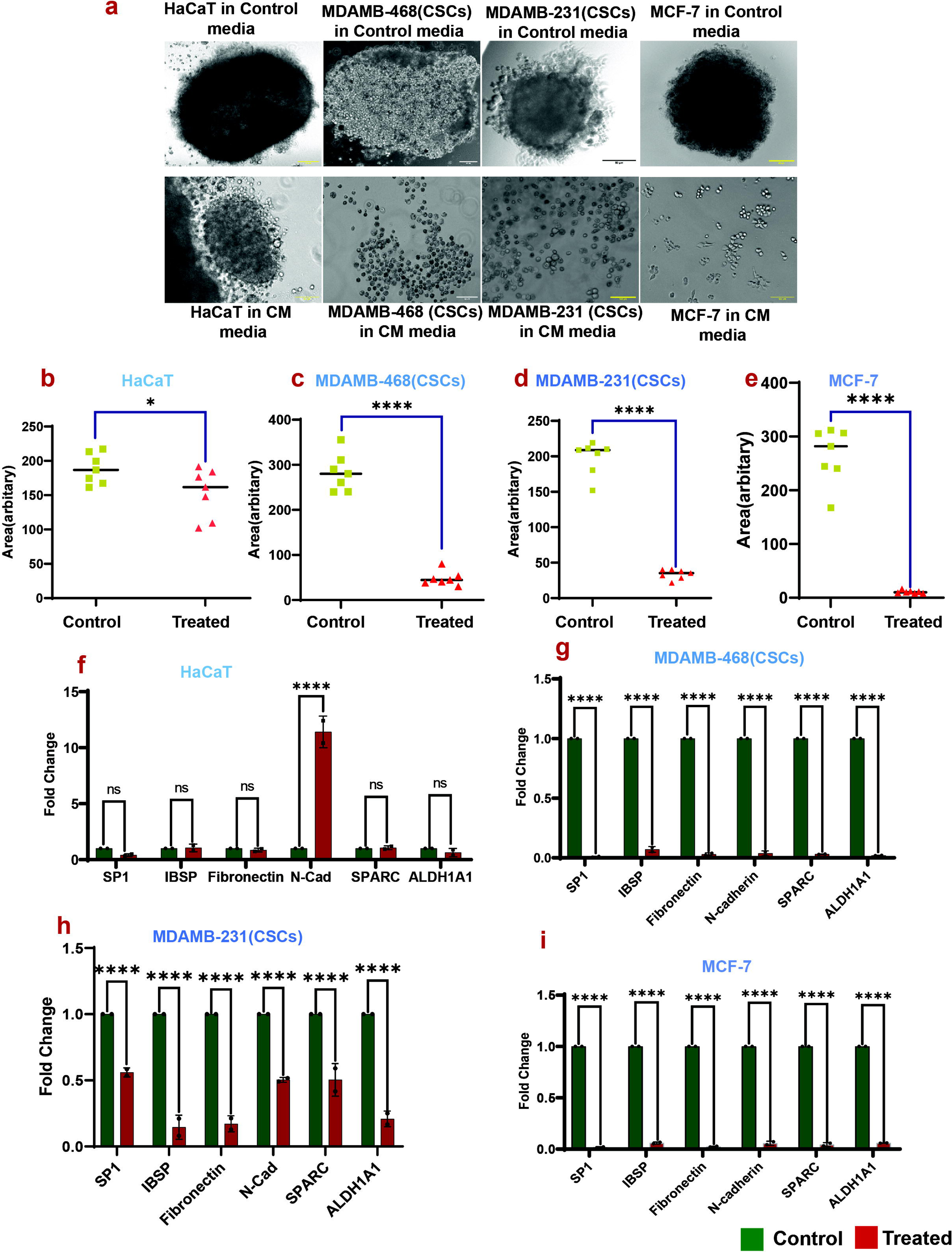
HSCs CM Disrupts TNBC CSCs and MCF-7 spheroid formation. **a)** 3D spheroid images of MDAMB-468(CSCs), MDAMB-231(CSCs), MCF-7, and HaCaT cells before and after CM treatment. **b-e)** The quantification of the spheroid area of CM-treated TNBC (CSCs), MCF-7, and HaCaT compared with untreated cells. **f-i)** gene expression of spheroid-derived CM-treated cells and control cells of TNBC CSCs, MCF-7 and HaCaT. Scalebar:100µm. Values are represented as Mean ± SEM. *p < 0.05, **p < 0.01, ***p < 0.001, ****p < 0.0001

## Discussion

This study unveils a novel therapeutic mechanism/possible intervention using hematopoietic stem cells (HSCs) against HER2^+^ and triple-negative breast cancer stem cells (TNBC-CSCs) by disrupting mitochondrial bioenergetics and inducing apoptosis. The HSCs’ tropism towards TNBC-CSCs is mediated by complex signaling and metabolic interactions, ultimately culminating in tumour cell death.

Initial characterization confirmed the successful isolation of HSCs and TNBC-CSCs from the cell lines. Next, co-culture trans-well assays demonstrated directed HSC migration towards TNBC-CSCs but not HaCaT cells, indicating preferential interaction of HSCs towards TNBC microenvironment **(Figure 1)**. Upon further probing for the molecular cues mediating this preferential migration followed by T cells differentiation of HSCs in the tumor microenvironment, we performed single-cell proteomics (SCP) of migrated HSCs. SCPs of migrated HSCs revealed altered expressions of several groups of proteins, namely cell adhesion proteins (LGALS1, ANXA2), cytoskeletal organization proteins (tropomyosin alpha 4 chains), stress response proteins (SOD1), and transcriptional regulation proteins (SMARCB1, HMGB1) **(Figure 2)**. Furthermore, we also observed the upregulation of IL-7 and Notch signaling components, suggesting their involvement in HSC-TNBC-CSC interactions. IL-7, known for promoting T-cell development (44), suggests potential HSC differentiation towards T-cell lineages within the tumor microenvironment. Flow cytometry confirmed the differentiation of migrated HSCs into CD4^+^ and CD8^+^ T cells, indicating a potential adaptive immune response. The role of these differentiated T-cells in tumor suppression requires further investigation.

Next, metabolomic analysis of HSC-conditioned media (CM) revealed enrichment in pathways including lactose synthesis, androgen/estrogen metabolism, selenoamino acid metabolism, pterine metabolism, amino sugar metabolism, fatty acid synthesis, galactose metabolism, sphingolipid metabolism, and tryptophan metabolism, indicating HSC derived metabolites enriched with the mentioned pathway (32). How do these metabolites influence TNBC-CSC metabolism? To learn more details, we performed global metabolomics of CM-treated TNBC-CSCs and MCF-7 cells. CM treatment of TNBC-CSCs and MCF7 cells induced distinct metabolic shifts, primarily affecting arginine biosynthesis, aromatic amino acid metabolism, branched-chain amino acid biosynthesis, pentose phosphate pathway, glutathione metabolism, histidine metabolism, and sphingolipid metabolism, suggesting paracrine effects disrupting TNBC-CSC metabolic homeostasis (45) **(Supplementary figure 6 & 7)**. For example, alterations in arginine metabolism could impact nitric oxide production, influencing tumor angiogenesis and immune evasion.

Next, we aimed to investigate the cell cycle regulation mechanism in CM-treated cells. Surprisingly, HSC-CM significantly altered the cell cycle dynamics of TNBC-CSCs by inducing G2/M arrest in MDAMB-468/231 (CSCs) and G0/G1 arrest in MCF7, correlating with the downregulation of CDKs 1, 2, 4, 5, 6, 7, and 9. The aforementioned results raise the next question: Do the differential cell cycle arrests reflect distinct cell cycle regulatory mechanisms in these cell lines? The more pronounced impact of HSC-CM on TNBC-CSCs hence indicates the potential therapeutic specificity of HSC-CM towards advanced TNBC **(Figure 3 & S7a)**.

As we have now demonstrated that HSC-CM substantially affects cell viability and cell cycle regulation in TNBC-CSCs, we next investigated the effects of HSC-CM on DNA damage and double-stranded DNA breaks. Results showed that CM treatment induced significant DNA double-strand breaks (DSBs), as evidenced by TUNEL and γ-H2AX staining, contributing to apoptosis. Cellular apoptosis was further corroborated by flow cytometry and analyses of pro-and anti-apoptotic gene and protein expression. Another interesting marker, the Forkhead box O3 protein status, was also checked and found to be activated in HSC-CM-treated CSCs. We then posed the question of whether FOXO3a activation was due to a direct consequence of DNA damage or a downstream effect of other signaling pathways activated by HSC-CM. The upregulation of FOXO3a, a regulator of apoptosis and DNA repair (46), may further support apoptosis induction. **(Figure 5)**

Next, we introduced the most important aspect of cell survival, namely cellular bioenergetics. The questions we discussed were: How did the shift in mitochondrial dynamics contribute to the observed metabolic changes and apoptosis? To find the answer, we examined the gene expression data of the major mitochondrial genes in the tested cell types following HSC-CM treatment. The key finding is the HSC-CM-mediated disruption of mitochondrial morphology and function. Evident mitochondrial fragmentation and perinuclear JC-1 aggregation indicated mitochondrial dysfunction. Altered expression of mitochondrial dynamics genes, with increased fission (Drp-1, Fis-1) and decreased fusion (Opa-1, Mfn-1/2), as well as changes in TOMM proteins, suggested disrupted mitochondrial homeostasis, potentially leading to increased ROS, DNA damage, and apoptosis (47). Altered expression of mitochondrial genes (SCO2, SURF1, COA6, MTCO1/2, COX4I1) further supports mitochondrial dysfunction. Again, this links to the question of how the HSC-CM-mediated alterations in mitochondrial dynamics induced metabolic changes and apoptosis in the TNBC-CSCs and the Her2+ MCF-7 cells. Critically, the aforementioned changes, such as mitochondrial gene expression induced by HSC-CM, provided a mechanistic link to the observed metabolomic alterations. For instance, changes in the expression of genes involved in oxidative phosphorylation (e.g., MTCO1/2, COX4I1) could directly impact cellular respiration and ATP production, potentially contributing to the observed shifts in central carbon metabolism pathways, such as the pentose phosphate pathway, which was altered in TNBC-CSCs treated with HSC-CM. Furthermore, disruption of mitochondrial dynamics and function can lead to increased production of reactive oxygen species (ROS), further influencing metabolic pathways and contributing to DNA damage and apoptosis (Figures 4 & 5). Specifically, the observed changes in sphingolipid metabolism in both HSC-CM and treated TNBC-CSCs could be linked to mitochondrial dysfunction, as sphingolipids are known to play a role in mitochondrial signaling and apoptosis (48) **(Figure 8)**.

Finally, HSC-CM disrupted TNBC-CSC spheroid formation, correlating with the downregulation of genes involved in spheroid formation, suggesting impaired self-renewal. Did HSC-CM target specific signaling pathways essential for stemness maintenance in TNBC-CSCs? Indeed, our gene expression analysis revealed significant downregulation of key genes involved in spheroid formation and stemness maintenance, including SP-1, IBSP, Fibronectin (38,39), N-cadherin (37,39), SPARC, and ALDH1A1 (37,41). SP-1 is a transcription factor that regulates the expression of several genes involved in cell growth, differentiation, and survival, including those involved in ECM production and cell adhesion (49). Downregulation of IBSP and Fibronectin, both extracellular matrix (ECM) components, suggested disrupted structural integrity of the spheroids, hindering their ability to maintain a cohesive 3D structure (35,36). N-cadherin, a cell adhesion molecule crucial for cell-cell interactions in spheroids, was also downregulated (37), further supporting the disruption of spheroid integrity. SPARC, an ECM-associated protein known to modulate cell-matrix interactions and inhibit cell adhesion, has been implicated in tumor suppression in some contexts (50,51). Its downregulation in our study might seem counterintuitive but could reflect a broader disruption of ECM homeostasis induced by HSC-CM. ALDH1A1, a marker of cancer stem cells, plays a crucial role in retinoic acid metabolism and has been associated with drug resistance and tumor initiation. Its downregulation suggests a direct impact of HSC-CM on the stem-like properties of TNBC-CSCs. This coordinated downregulation of genes essential for spheroid formation and stemness strongly suggests that HSC-CM effectively targets and impairs the self-renewal capacity of TNBC-CSCs **(Figure 7)**.

In conclusion, this study provides compelling evidence that HSCs may be a promising therapeutic strategy for TNBC-CSCs. HSCs exhibit targeted tropism mediated by complex signaling and metabolic crosstalk. HSC-CM induces metabolic stress, cell cycle arrest, DNA damage, apoptosis, and disruption of mitochondrial dynamics/function in TNBC-CSCs. These findings warrant further investigation into HSC-derived factors and preclinical evaluation of this therapeutic approach. Moreover, for clinical setup, HSCs must be cultured from clinical samples using xeno-free media conditions, which may also have novel implications for clinical translation.

## Ethical Statement

Ethics was approved by Yenepoya Ethics Committee YEC-1/2022/219 and YEC2/2024/006, Yenepoya Medical College, Yenepoya (Deemed to be University).

## CRediT authorship

SM conceptualized, designed, and performed all the experiments, analyzed the data, and wrote the initial manuscript. SB acquired the funding, provided critical comments and edited the manuscript. BB conceptualized the project, supervised and designed the experiments, and edited the manuscript. SS designed the experiments, edited the result figures, and revised the manuscript. All authors approved the final version of the manuscript.

## Acknowledgement

The authors thank Mr. Muhammad Nihad, SRF, SCRMC, for helping with cell sorting. We are also thankful to Dr. Zuzana Kečkéšová, Institute of Organic Chemistry and Biochemistry, Czech Academy of Sciences, Prague, Czech Republic, for critical suggestions in experimental design. We also thank the Yenepoya Research Centre for its instrumental facilities.

## Availability of Data and Materials

The mass spectrometry proteomics data have been deposited in the ProteomeXchange Consortium via the PRIDE partner repository, with the dataset identifier **PXD061260**. Metabolomics data have been submitted to the Massive Data Repository using the dataset identifier **MSV000097239**. Other data generated or analyzed during this study are included in the article and its supplementary information files or are available from the corresponding author upon reasonable request.

